# Disentangling contact and ensemble epistasis in a riboswitch

**DOI:** 10.1101/2022.10.27.514099

**Authors:** Daria R. Wonderlick, Julia R. Widom, Michael J. Harms

## Abstract

Mutations introduced into macromolecules often exhibit epistasis, where the effect of one mutation alters the effect of another. Knowledge of the mechanisms that lead to epistasis is important for understanding how macromolecules work and evolve, as well as for effective macromolecular engineering. Here we investigate the interplay between “contact epistasis” (epistasis arising from physical interactions between mutated residues) and “ensemble epistasis” (epistasis that occurs when a mutation redistributes the conformational ensemble of a macromolecule, thus changing the effect of the second mutation). We argue that the two mechanisms can be distinguished in allosteric macromolecules by measuring epistasis at differing allosteric effector concentrations. Contacts give rise to epistasis in the microscopic equilibrium constants describing the conformational ensemble. Ensemble epistasis manifests in thermodynamic observables, such as the energy of ligand binding or enzyme activation, that depend on the concentration of allosteric effector. Using this framework, we experimentally investigated the origins of epistasis in three mutant cycles introduced into the adenine riboswitch aptamer domain. We found evidence for both contact and ensemble epistasis in all cycles. Further, we found that the two mechanisms of epistasis can interact with each other. For example, in one mutant cycle we observe contact epistasis of 6 kcal/mol attenuated by the ensemble to only 1.5 kcal/mol in the final thermodynamic observable. Finally, our work yields simple heuristics for identifying contact and ensemble epistasis using limited experimental measurements.

**Statement of significance:** Mutations to protein or RNA molecules often have different effects when introduced individually versus together. To understand and engineer biological macromolecules, we must identify the mechanistic origins of this phenomenon. Here, we measured the interplay between direct, physical interactions between mutations (“contact epistasis”) and indirect interactions mediated by conformational ensembles (“ensemble epistasis”). We introduced pairs of mutations into an RNA molecule that transitions between several different conformations. We found epistasis arising from both contacts and the ensemble, and that the two mechanisms could synergize with one another. Our work reveals that one must consider the effects of mutations on multiple conformations to understand epistasis and suggests a few rules-of-thumb for disentangling contact and ensemble epistasis in other macromolecules.

## Introduction

Mutations often exhibit epistasis, where the effect of one mutation alters the effect of another (1, 2). Intramolecular epistasis—interactions between mutations within the same macromolecule—is useful for dissecting molecular function (3, 4), can shape molecular evolution (2, 5–7), and must be properly captured for macromolecular engineering (8–10). Knowledge of the mechanisms of epistasis is thus important for both understanding and designing macromolecules.

Broadly, there are two sources of epistasis in macromolecules. The most intuitive is mediated by structural contacts between the sites. Such contacts could be simple one-to-one interactions (e.g. ion pairs, base pairs, hydrogen bonds, etc.) (7, 11, 12) or larger networks of interactions (e.g. pathways of residues connecting distant sites) (4, 13). The second mechanism arises from the ensemble of conformations populated by the macromolecule (14–16). This epistasis arises when a macromolecule populates several conformations that respond differently to the same mutation. A mutation at one site redistributes the relative populations of the conformations, thus altering the effect of a second mutation. The relative contributions of contact and ensemble mechanisms to epistasis across macromolecules are unknown.

We recently found that environment-dependence is a diagnostic feature of ensemble epistasis (14). If one changes the environment such that the distribution of conformations is altered, one can change the contribution of ensemble epistasis. Concentration-dependent epistasis is relatively simple to observe and interpret for an allosteric macromolecule, where altering the concentration of an allosteric effector redistributes the ensemble—and thus the measurable output—of the system in a predictable fashion. We recently demonstrated ensemble epistasis experimentally using the lac repressor, which exhibited epistasis in DNA binding as a function of its allosteric effector IPTG (17). Intriguingly, we also found that redistribution of the ensemble could not fully explain the epistasis we observed—suggesting that other mechanisms besides ensemble epistasis were at play in the protein.

Our previous work raised several questions. How can we distinguish between epistasis arising from contacts, the ensemble, or both simultaneously? What are the relative magnitudes of these two forms of epistasis? How do they influence one another? Is ensemble epistasis within the purview of proteins alone, or does it arise in other classes of macromolecules?

To answer these questions, we investigated the contributions of contact and ensemble epistasis between pairs of mutations in the well-characterized adenine riboswitch aptamer domain. This RNA molecule has a three-state conformational ensemble that allows it to bind to adenine in a magnesium-dependent fashion (18–25). We measured magnesium-dependent epistasis between mutations we expected to have varying contributions of contact and ensemble epistasis, finding evidence for both mechanisms in all mutant cycles. Our work reveals that contact and ensemble epistasis can have comparable magnitudes, and that the two sources of epistasis can interact in highly nonlinear ways. Further, our results suggest that one can use environment-dependence as a useful heuristic for disentangling ensemble and contact epistasis in macromolecules.

## Materials and Methods

### RNA sample preparation

We synthesized sequence variants of the *V. vulnificus* adenine riboswitch aptamer domain by *in vitro* transcription of the corresponding DNA oligonucleotides (HiScribe T7 High Yield RNA Synthesis Kit, New England Biolabs; oligonucleotides ordered from Eurofins). The wildtype RNA sequence is shown with mutated positions in boldface: 3’GGGAAGAUAUAAUCCUAAUGAU**A**UG**G**UUUGGGAGUUU**C**UACCAAGAG**C**CUUAA ACUCUUGAUUAUCUUCCC. The riboswitch product was purified by running the *in vitro* transcription reaction on a 12% denaturing polyacrylamide gel, identifying the 71-nt product by UV shadowing, and extracting the product from the gel by electroelution. The purified RNA product was concentrated by ethanol precipitation, desalted, quantified by Nanodrop absorbance spectroscopy, and resuspended in 50 mM Tris-HCl before being stored at −80 °C until further use.

### Measurement of 2-aminopurine binding

We measured binding of 2-aminopurine (2AP), a fluorescent analog of adenine, to the riboswitch aptamer domain using a previously published protocol (19, 25). Riboswitch aliquots were thermally denatured at 90 °C for 1 min before refolding on ice for approximately 10 min. Increasing concentrations of RNA (0.125-8 μM) were combined with 50 nM 2AP, 100 mM KCl, and 0.1-100 mM MgCl2 in 96-well black bottom plate. The assay plate was shaken in a Molecular Devices SpectraMax i3 fluorescence plate reader at 37 °C for 5 minutes to allow the reaction mixtures to reach equilibrium. 2AP exhibits high fluorescence in water and quenched fluorescence when bound to the riboswitch. We excited each sample at 310 nm and measured emission between 335 and 450 nm. We integrated each emission spectrum and converted the area to fraction of 2AP bound using a negative control (0 nM RNA, 0 nM 2AP) and a positive control (0 nM RNA, 50 nM 2AP). Because 2AP fluorescence was negligible in samples where the fluorophore was completely bound by RNA, we used a negative control lacking 2AP entirely to universally mimic this fully quenched state. We averaged two technical replicates for each condition per experiment and collected at least three biological replicates for each experimental condition.

### RNA ensemble modeling

As described in the results, we used a previously validated thermodynamic model of the riboswitch ensemble to extract information about the populated conformations (23). This model describes the riboswitch with two apo conformations—extended (E) and docked (D)—each of which can form a complex with 2AP (E·A and D·A). We tested three variants of the model: a two-state model with conformations E and D·A, a three-state model with conformations E, D, and D·A, and a four-state model with conformations E, E·A, D, and D·A. We also tried subvariants of these models in which we fixed the values of the different equilibrium constants. As described in the results, we found that the three-state model with Kdock set to 1.0 was able to reproduce our binding data without overfitting. A mathematical description of the three-state model follows; the two-state and four-state models are given in the supplement (Fig S1).

The three-state model is shown schematically in Fig 1A. The concentrations of the E, D, and D·A conformations are given by:

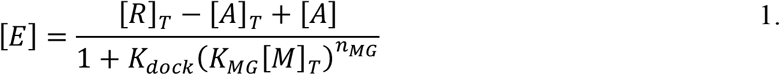

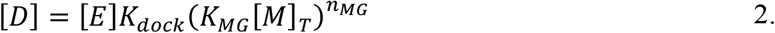

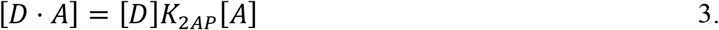

where K_dock_ is the equilibrium between E and D in the absence of Mg^2+^ or 2AP, K_2AP_ is the affinity of 2AP for the D conformation, K_MG_ is the relative affinity of the D and E conformations for Mg^2+^, nMG is the difference in the number of Mg^2+^ ions bound by the D and E conformations, [R]_T_ is the total concentration of RNA, [M]_T_ is the total concentration of Mg^2+^, [A]_T_ is the total concentration of 2AP, and [A] is the concentration of free 2AP. Our experimental observable, the fraction of 2AP bound to RNA, is given by:

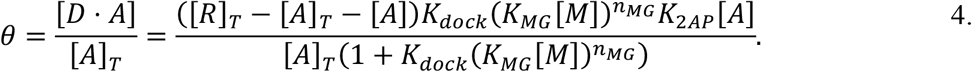

**Figure 1.**
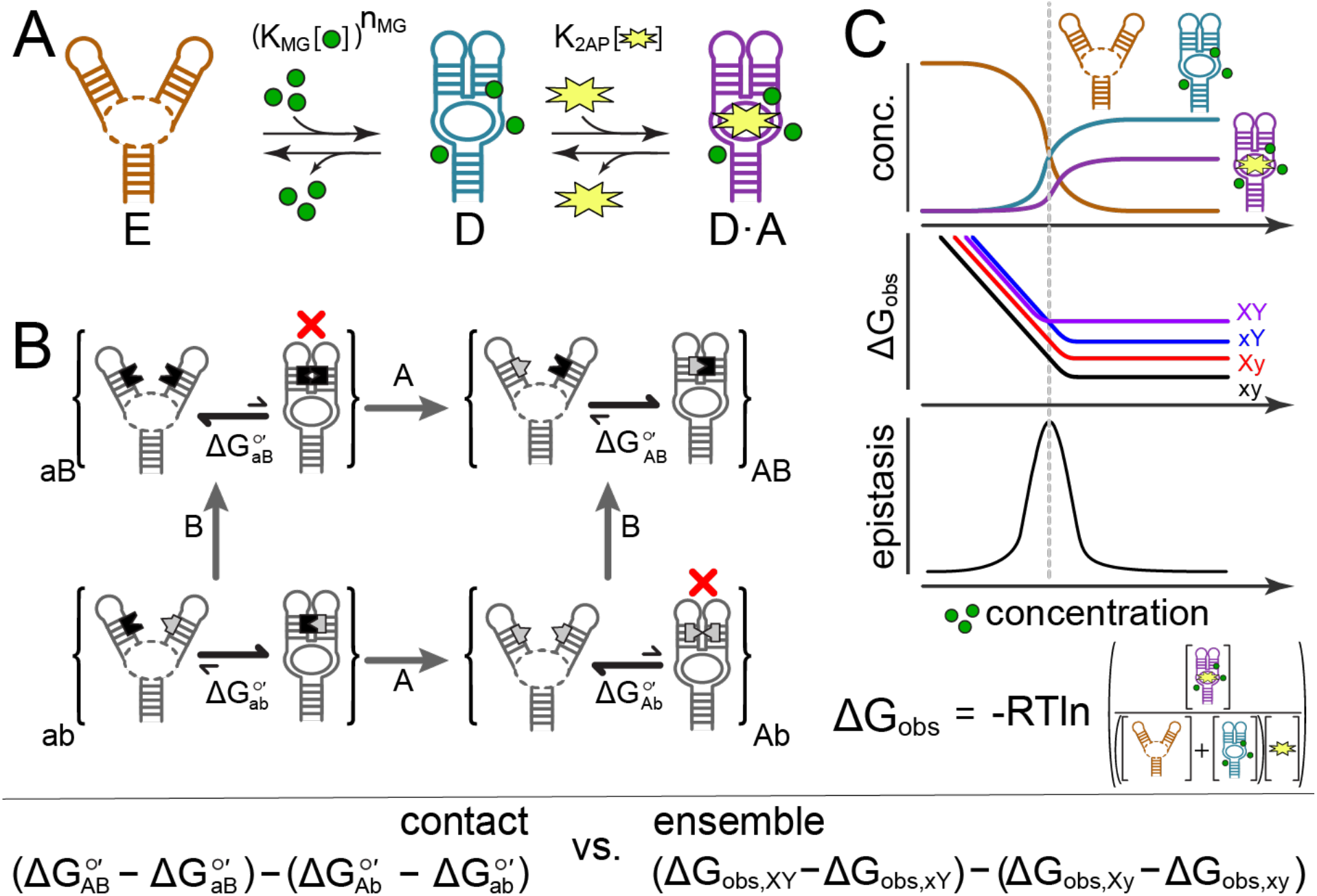
Differentiating contact and ensemble epistasis in the adenine riboswitch. A) A thermodynamic model of the riboswitch ensemble. The molecule can be extended (E), docked (D), or docked and bound to 2AP (D·A). The equilibria are defined by three parameters: K_MG_, n_MG_, and K_2AP_. The equilibria are tuned by the concentration of Mg^2+^ ions (green circles) and 2AP (yellow star). B) Epistasis arising from a contact between two sites. RNA molecules within braces represent an ensemble for a specific genotype (ab, Ab, aB, or AB). The equilibrium between the E and D forms is shown by arrows within the braces and measured by ΔG°’. The A and B mutations are indicated by bold gray arrows between ensembles. Bases at these sites can either be indented (black) or protruding (gray). Mutation A flips the left base from indented to protruding; mutation B flips the right base from protruding to indented. Mismatches are indicated with a red “×”. C) Epistasis arising from the ensemble. The cartoon graphs show concentrations of E, D, and D·A for wildtype (top); ΔG_obs_ for four genotypes (xy, Xy, xY, and XY) taken from a hypothetical mutant cycle (middle); and epistasis in ΔG_obs_ calculated from the middle plot (bottom). These series are plotted as a function of Mg^2+^ concentration. The mathematical expressions for contact and ensemble epistasis are displayed at the bottom of the figure.

This model has four explicit parameters: K_dock_, K_MG_, nMG, and K2AP. It also has two implicit parameters: [M] (the concentration of free Mg^2+^) and [A] (the concentration of free 2AP).

We implemented this model as a function that returns the concentrations of all relevant species ([A], [M], [E], [D], and [D·A]) given the values of the thermodynamic parameters (K_dock_, K_2AP_, K_MG_, and n_MG_) and total species concentrations ([R]_T_, [M]_T_, and [A]_T_). This function encodes the thermodynamic relationships above (equations 1–4) and enforces mass-balance relationships (e.g. [A]_T_= [A] + [D·A]). Internally, this function guesses values for [A] and then iterates to self-consistency between the thermodynamic and mass-balance relationships.

To estimate the values of the thermodynamic parameters consistent with our binding data, we used a Bayesian Markov-Chain Monte Carlo (MCMC) strategy to sample over parameter combinations. We analyzed all experimental conditions simultaneously for each genotype, globally estimating the thermodynamic parameters. We used the following likelihood function:

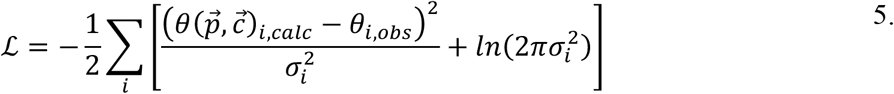

where θ_i,obs_ and σ_i_ are the mean and standard deviation of the measured 2AP fractional saturation at condition *i*; θ_i,caic_ is the calculated 2AP fractional saturation under these conditions given a vector of parameters 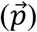 and total concentrations 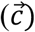. If our experimental σ_i_ was < 0.05, we set its value to 0.05, thus enforcing a minimum uncertainty on the mean of any observed data point. We performed MCMC sampling on the natural logs of all equilibrium constants. We generated 15,000 MCMC samples from 100 MCMC walkers for each genotype, discarding the first 1,000 as burn in. We checked for convergence by comparing results from multiple independent runs. For the final three-state, three-parameter model we used in our analysis, we generated a million MCMC samples from six different sampling runs. We used the emcee 3.1.0 (26) and the “likelihood” python libraries (https://github.com/harmslab/likelihood) for these calculations. We implemented the riboswitch binding models in Python 3.9 extended with numpy 1.21.1 (27), pandas 1.3.1 (28), and scipy 1.6.2 (29).

### Data availability

All experimental data, software, and scripts to reproduce analyses reported in this manuscript are available on github (https://github.com/harmslab/riboswitch-epistasis).

## Results

### Selecting the adenine riboswitch as a model system

We set out to measure the relative contributions of contact and ensemble epistasis using mutant cycles introduced into the adenine riboswitch aptamer domain, a 71-nucleotide RNA molecule composed of three helices surrounding an adenine binding pocket (30) (Fig 2A). We selected this RNA molecule as our model system because it has a relatively simple, well-defined conformational ensemble consisting of two conformations: an extended form (E) and a compact, docked-loop form (D) (Fig 1A). Relative to the E conformation, the D conformation has extensive base pairs, hydrogen bonds, and hydrophobic packing between the two helical arms of the riboswitch. This forms an adenine binding pocket unique to the D conformation; therefore, the D conformation has much higher affinity for adenine than the E conformation. Mg^2+^ ions are key allosteric effectors for the riboswitch ensemble because they nonspecifically neutralize repulsion between backbone phosphate groups to promote formation of the compact D conformation, thus promoting adenine binding (23, 31, 32).

**Fig 2.**
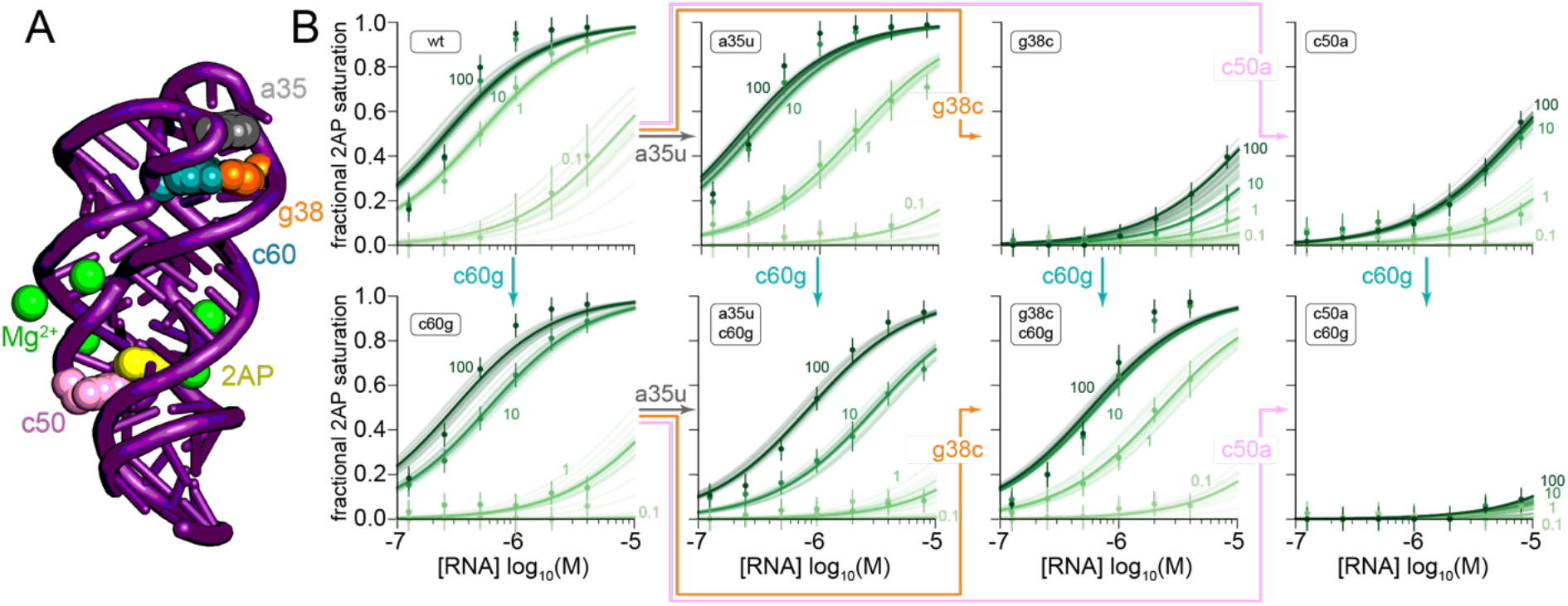
2AP binding behavior can be captured by a thermodynamic model of the riboswitch ensemble. A) Crystal structure of the adenine riboswitch aptamer domain (PDB ID: 5SWE) (35). The overall architecture is shown as a purple cartoon. Bases mutated in this study are indicated with colored spheres (labeled on structure). The bound adenine base is shown as yellow spheres; probable Mg^2+^ ion locations are shown as green spheres, taken from PDB: 1Y26 (36). B) Each panel shows 2AP binding for the genotype indicated. Columns are (left to right) wildtype, a35u, g38c, and c50a. Rows are without (top) and with (bottom) the c60g mutation. The arrows between panels define three mutant cycles: a35u/c60g, g38c/c60g, and c50a/c60g. Each panel shows the fraction of 2AP bound, measured by relative 2AP fluorescence compared to free and fully quenched controls, as a function of riboswitch concentration. The green shade indicates the total Mg^2+^ concentration, labeled on each plot in millimolar. Each point is the average of at least three experimental replicates, with standard deviations indicated as error bars. Lines were calculated using parameters taken from 50 randomly selected MCMC samples of the three-state/three-parameter model shown in Fig 1A.

Quantifying epistasis between mutations to the riboswitch requires measuring how mutations affect a biological function when introduced individually versus together. As a proxy for the biological function of the riboswitch, we measured its binding to 2-aminopurine (2AP), a fluorescent analog of adenine. The 2AP binding energy is given by:

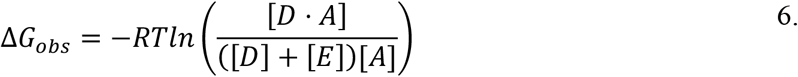

where the conformations are those labeled in Fig 1A. We can calculate the concentrations of each of these species using Equations 1–3, which require knowing the total concentrations of the molecular components (Mg^2+^, RNA, and 2AP), two apparent equilibrium constants (K_MG_ and K_2AP_), and an ion uptake coefficient (n_MG_). We can rewrite our 2AP binding energy in terms of these equilibrium constants by:

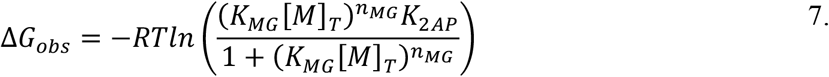

where [M]_T_ is the total Mg^2+^ concentration. The 2AP binding energy can also be calculated directly from experimental measurements of the fraction of 2AP bound to RNA using:

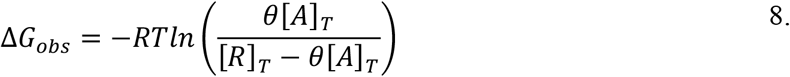

where [A]_T_ is the total 2AP concentration, [R]_T_ is the total RNA concentration, and θ is the fraction of 2AP bound to RNA (Equation 4). Describing ΔG_obs_ in terms of the system equilibrium constants (Equation 7) or fraction of 2AP bound to RNA (Equation 8) allows us to calculate a single thermodynamic observable that encapsulates the riboswitch’s conformational ensemble without needing to directly measure the population of each conformation spectroscopically.

### Contact and ensemble epistasis will give different signals

We next asked how contact and ensemble epistasis would manifest conceptually and mathematically within the riboswitch system. Contact epistasis occurs when two mutations interact within a conformation, thus changing the energy of that conformation. Consider the predicted effects of a pair of mutations (a→A and b→B) at residues that form complementary contacts in the docked conformation (D) of the riboswitch (Fig 1B). When introduced individually, these mutations create non-complementary interactions and shift the equilibrium towards the extended form (E). When introduced together, however, complementarity is restored, thus pushing the equilibrium back towards the docked conformation. Such contact epistasis will manifest in the microscopic equilibrium between these two conformations. This can be quantified using the biological standard state free energy for this equilibrium, ΔG°’ =-RTln(K_eq_), yielding the following expression for contact epistasis:

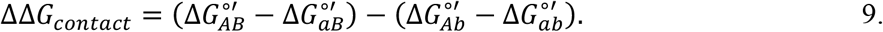

Contact epistasis is independent of the concentrations of other conformations in solution–such as that of Mg^2+^—because it arises from interactions within a conformation rather than a non-linear redistribution of conformations. Thermodynamically, contact epistasis occurs in standard state free energies to reflect that the interactions are concentration independent.

Ensemble epistasis, by contrast, occurs when a mutation redistributes the relative populations within the ensemble, thus changing the effect of a second mutation on an ensemble-averaged observable. Consider another pair of mutations (x→X and y→Y) to the riboswitch (Fig 1C). If each mutation changes one or more of the equilibrium constants (Equation 7), each of the xy, Xy, xY and XY riboswitch variants will have different populations of the E, D, and D·A conformations at a given Mg^2+^ concentration. The altered equilibrium constants change the apparent 2AP affinity (ΔG_obs_) for each variant in a nonlinear fashion. One can calculate ensemble epistasis as:

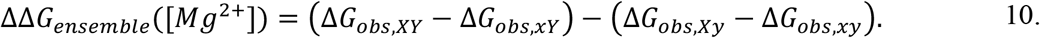

Ensemble epistasis depends on the Mg^2+^ concentration because the population of the ensemble– and thus ΔG_obs_—is controlled by [Mg^2+^]. At low Mg^2+^ concentration, only the E conformation is populated. At high Mg^2+^ concentration, we expect ΔG_obs_ to plateau because 2AP becomes limiting. Additional Mg^2+^ can no longer change the relative population of D·A and thus can no longer alter ΔG_obs_. At intermediate Mg^2+^ concentrations, multiple conformations are populated, thus maximizing the redistributive effects of each mutation and the magnitude of ensemble epistasis (shown schematically in Fig 1C).

In summary, contact epistasis appears in the standard state free energy of a specific conformation, while ensemble epistasis appears in an ensemble-averaged observable, such as a free energy of 2AP binding. Because contact epistasis could, in principle, strongly perturb the population of the conformational ensemble, we expect a complicated interplay between the two mechanisms. Without experimental knowledge of the relative magnitudes of each mechanism, however, we cannot predict what this synergy might look like for a real macromolecule.

### Mutant cycle design

We next designed three mutant cycles for which we predicted different contributions of contact and ensemble epistasis. We built these cycles from four mutations that were previously reported to alter the riboswitch’s 2AP and Mg^2+^ affinity (Fig 2A) (19, 20). All cycles involve c60g (Fig 2A; blue), paired with either a35u, g38c, or c50a (Fig 2A; gray, orange, and pink). For consistency, we measured epistasis across all cycles as the difference between the effect of c60g introduced into a mutant riboswitch versus the wildtype riboswitch. In the wildtype background, c60g disrupts the complementarity of a Watson-Crick-Franklin base pair at the docked-loop interface of the D conformation, leading to a decrease in 2AP affinity (19, 20).

We expected the a35u/c60g pair to exhibit minimal epistasis of either type. a35u has been reported to slightly increase ligand binding affinity by adding an extra hydrogen bond to the docked-loop interface (20). a35 is not in direct contact with c60, and the two sites are not in allosterically linked regions of the riboswitch (Fig 2A). We therefore anticipated little epistasis between these positions.

We expected the g38c/c60g pair to give strong contact epistasis. g38 and c60 form a Watson-Crick-Franklin base pair at the docked-loop interface of the wildtype riboswitch’s D conformation (Fig 2A) (19, 20). When introduced individually, these mutations disrupt this base pair and thus greatly decrease 2AP binding affinity. When both mutations are introduced together, c38/g60 form a reversed Watson-Crick-Franklin base pair, which has been shown to restore 2AP binding activity (19).

We expected the c50a/c60g pair to exhibit significant ensemble epistasis but little contact epistasis. Positions 50 and 60 are ~40 Å apart in the crystal structure (Fig 2A). c60 is at the docked-loop interface, while c50 is directly adjacent to the 2AP binding pocket. The c50a mutation has been shown previously to reduce 2AP binding affinity by destabilizing the ligand binding pocket (20). Because the docked-loop region and 2AP binding pocket are in allosteric communication, we anticipated epistasis between these two mutations; however, we predicted this would be mediated by the ensemble rather than a direct contact given the separation between the positions.

### Defining the thermodynamic ensemble

With our system selected and our mutant cycles designed, we next set out to measure how the mutations perturbed the riboswitch ensemble. Previously, Leipply and Draper validated a thermodynamic model with two apo conformations of the adenine riboswitch—extended (E) and docked (D)—each of which can bind 2AP (E·A and D·A) (23). In a separate study (31), they found they could model RNA-Mg^2+^ interactions with an ion uptake coefficient that captured the number of excess Mg^2+^ ions bound by D relative to E. Put together, this results in a five parameter thermodynamic model (Fig S1) with the following terms: K_dock_ (the equilibrium between E and D in the absence of Mg^2+^), K_2AP_ (the affinity of the riboswitch for 2AP), K_link_ (the coupling between Mg^2+^/2AP binding and formation the D conformation), K_MG_ (the Mg^2+^ affinity of the D conformation relative to the E conformation), and n_MG_ (the difference in the number of Mg^2+^ ions bound by the D and E conformations).

To resolve the parameters in this model, we measured 2AP binding over 4 orders of magnitude in Mg^2+^ concentration: 0.1, 1, 10, and 100 mM. This covers the physiological range of Mg^2+^ concentrations (33). We varied RNA concentrations over ~2 orders of magnitude, from 0.125 to 8 μM, while maintaining a constant, limiting 2AP concentration of 50 nM. The results of these experiments for all eight constructs are shown in Fig 2B. As expected, we observed increasing 2AP binding with increasing amounts of RNA for all variants. Further, we observed a link between 2AP binding and Mg^2+^ concentration, with the midpoint of the 2AP binding curves decreasing with increasing Mg^2+^concentration.

We used a Bayesian Markov Chain Monte Carlo (MCMC) strategy to sample model parameters for each riboswitch variant that could reproduce our experimental observations. The four-state/five-parameter model reproduced our measured binding data well (Fig S2). We noted, however, that the E·A conformation never accounted for more than 0.02% of the total RNA concentration for all datasets. We therefore asked if we could simplify our model by ignoring the E·A conformation, thus eliminating the model parameter K_link_. This three-state/four-parameter model reproduced our experimental data as well as the original model (Fig S2). We also noticed strong covariation between K_MG_ and K_dock_ across MCMC samples, so we tried a model in which we set K_dock_ to 1.0. In this formulation, K_MG_ is an apparent constant that accounts for both the energy of Mg^2+^ binding and the relative stability of the D versus E conformations. This three-state/three-parameter model still reproduced our experimental data well (Fig 2B).

Finally, we asked if we could simplify the model further by describing the system with only two conformations, E and D·A. This two-state model makes the riboswitch cooperative, meaning Mg^2+^ binding necessitates 2AP binding. This two-state model could not reproduce our measured 2AP binding data (Fig S2). We used an Akaike Information Criterion (AIC) test to validate our visual inspection of the binding curves. The AIC test favors models with high likelihoods while penalizing models with excess parameters (34). Our AIC test favored the three-state/three-parameter model (Table S1). All fit parameters for the final three-state/three-parameter model are given in Table S2.

### Mutant Cycle #1: a35u/c60g

We next analyzed the epistasis within each mutant cycle, starting with a35u/c60g. These bases do not contact on another and are distant from both the Mg^2+^ and 2AP binding sites (Fig 2A). We therefore expected relatively weak contact and ensemble epistasis for this mutant pair.

We first looked for contact epistasis in the a35u/c60g mutant cycle. We extracted marginal distributions for the values of ΔG°’_2AP_, ΔG°’_MG_ and n_MG_ from the MCMC samples for the relevant variants (Fig 3A-C) and calculated epistasis in each parameter using Equation 9. c60g had no detectable effect on ΔG°’_2AP_ when introduced alone, but increased ΔG°’_2AP_ by 0.6 [0.4, 0.8] kcal/mol in the a35u background. (In this notation, we write the median effect followed by the 95% credibility interval extracted from the marginal distribution in brackets). If the mutations exhibited no epistasis, we would expect the a35u/g38c double mutant to have ΔG°’_2AP_corresponding to the green point on Fig 3A. The actual change is larger, reflecting measurable epistasis in ΔG°’_2AP_ (0.52 [0.28, 0.73]). We saw little evidence of epistasis in ΔG°’_MG_ or n_MG_: a35u and c60g both increased ΔG°’_MG_ by ~0.5 kcal/mol (Fig 3B) and neither mutation had a measurable effect on n_MG_ (Fig 3C)

**Figure 3.**
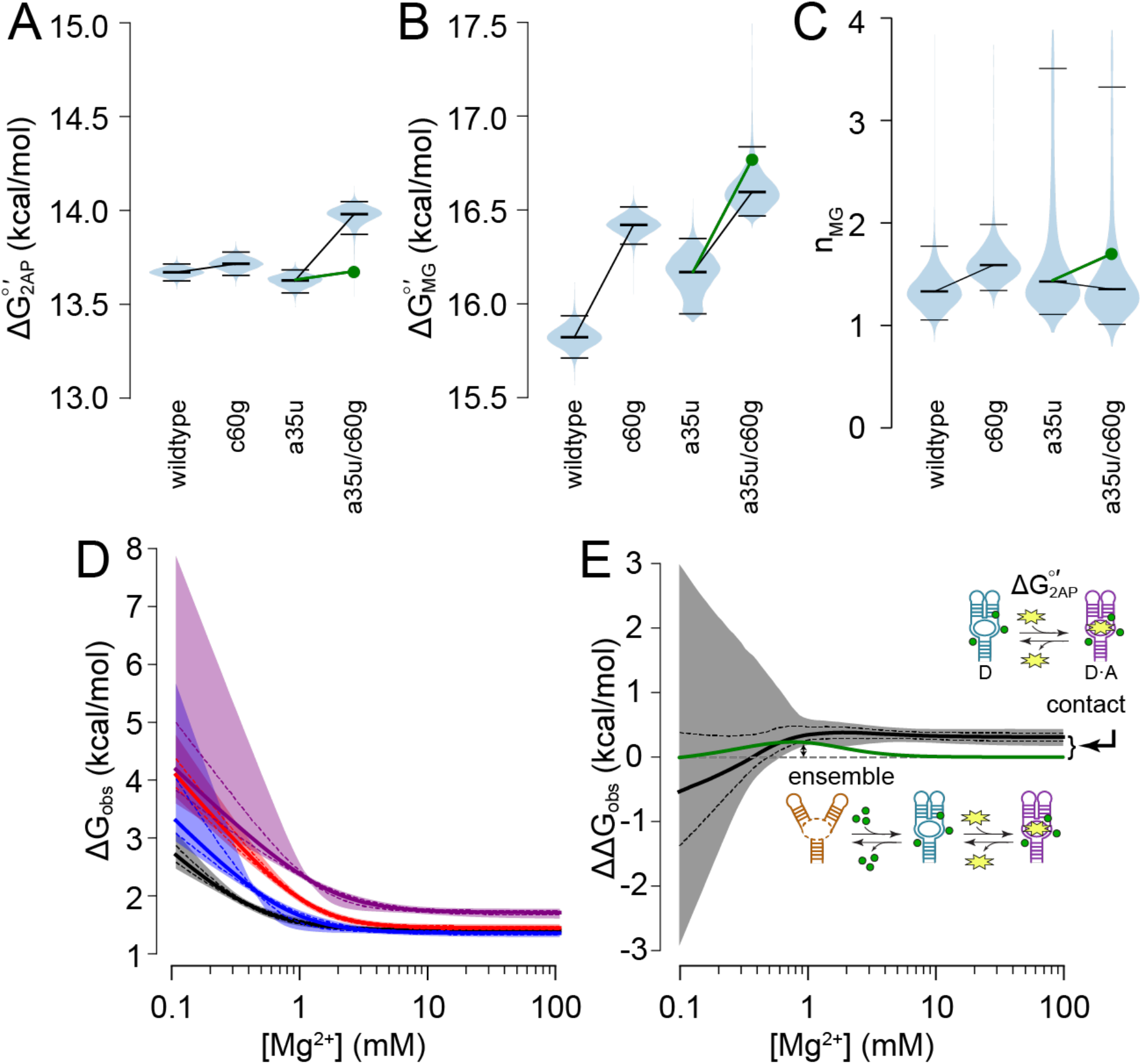
Contact and ensemble epistasis in the a35u/c60g cycle. Panels A-C show marginal distributions of ΔG°’_2AP_, ΔG°’_MG_, and n_MG_ taken from across all MCMC samples. The area of the violin plot encompasses all values seen. The central bar is the median; the top and bottom bars mark the 95% credibility interval. The black lines between series indicate the median effect of the c60g mutation in the wildtype (left) or a35u (right) backgrounds. Epistasis appears as a background-dependent difference in the slope of the line; the green lines and points indicate the expected value if the mutations do not exhibit epistasis. D) ΔG_obs_ versus Mg^2+^ concentration calculated using Equation 7 for wildtype (black), c60g (red), a35u (blue) and a35u/c60g (purple). Solid lines indicate the median of the MCMC samples; dashed lines indicate the standard deviation; areas indicate the 95% credibility region. E) Epistasis in ΔG_obs_ calculated from the curves in panel D. The green line indicates ΔG_obs_ calculated in the absence of contact epistasis (green points in panels A-C). The small arrows with cartoons indicate the Mg^2+^ concentrations where epistasis arises from the ensemble (1 mM) versus contacts (>10 mM).

We next looked for evidence of ensemble epistasis in the a35u/c60g mutant cycle. We used Equation 7 to calculate ΔG_obs_ versus Mg^2+^ concentration for all four variants (Fig 3D). Each variant gave decreasing ΔG_obs_ with increasing Mg^2+^ concentration, reflecting the shift in equilibrium from E to D·A (Fig S3). ΔG_obs_ plateaued for all variants when the concentration of 2AP became limiting: additional Mg^2+^ could no longer change the relative concentration of D·A and thus could no longer alter ΔG_obs_. We then calculated epistasis in ΔG_obs_ using Equation 10 (Fig 3E). At low Mg^2+^concentration, our uncertainty was high and the signal indistinguishable from zero; for Mg^2+^ above 1 mM, we observed approximately constant epistasis in ΔG_obs_ (0.31 [0.17,0.43] kcal/mol).

We next investigated the interplay between contact and ensemble epistasis in the a35u/c60g mutant cycle by calculating magnesium-dependent epistasis in ΔG_obs_ after removing contact epistasis. We assumed that the parameter values for a35u/c60g double mutant were given by the sum of the median effects of a35u and c60g introduced alone, thereby eliminating epistasis in ΔG°’_2AP_, ΔG°’_MG_ and n_MG_. These non-epistatic parameters are shown as the green points for a35u/c60g in Fig 3A-C. The epistasis in ΔG_obs_ we observe in the absence of contact epistasis is shown as a green line in Fig 3E. The curve behaves as expected for a system exhibiting ensemble epistasis. It peaks when the diversity of the ensemble is maximized and tails off to zero when the system populates a single conformation at high and low Mg^2+^ concentrations (Fig S3). The ensemble contributes maximally to the epistasis in this mutant cycle around 1 mM Mg^2+^ (labeled arrow, Fig 3E). At higher Mg^2+^ concentrations, contact epistasis in 2AP binding (ΔG°’_2AP_) is the source of epistasis between the mutations (labeled arrow, Fig 3E).

### Mutant Cycle #2: g38c/c60g

We next turned to the g38c/c60g mutant cycle. We expected these mutations to yield high-magnitude contact epistasis because the nucleotides form a Watson-Crick-Franklin base pair in the D conformation that inverts complementarity over the mutant cycle (Fig 2A).

As with the previous mutant cycle, we looked for evidence of contact epistasis in the g38c/c60g mutant cycle by considering the standard state free energies for the equilibria defining the system. We calculated the marginal distributions of ΔG°’_2AP_, ΔG°’_MG_ and n_MG_ for all four variants (Fig 4A-C). The parameters were well constrained for the wildtype, c60g, and g38c/c60g variants, but not g38c introduced alone. The g38c variant exhibits low 2AP binding, even at high Mg^2+^ and RNA concentrations (Fig 2B). Based on our MCMC samples, we are confident that g38c differs from the other variants; however, this could arise because the mutation increases ΔG°’_MG_, increases ΔG°’2AP, or some combination of the two (Fig 4D). This leads to a strong inverse correlation in estimates of ΔG°’_2AP_ and ΔG°’_MG_ (Fig 4D).

**Figure 4.**
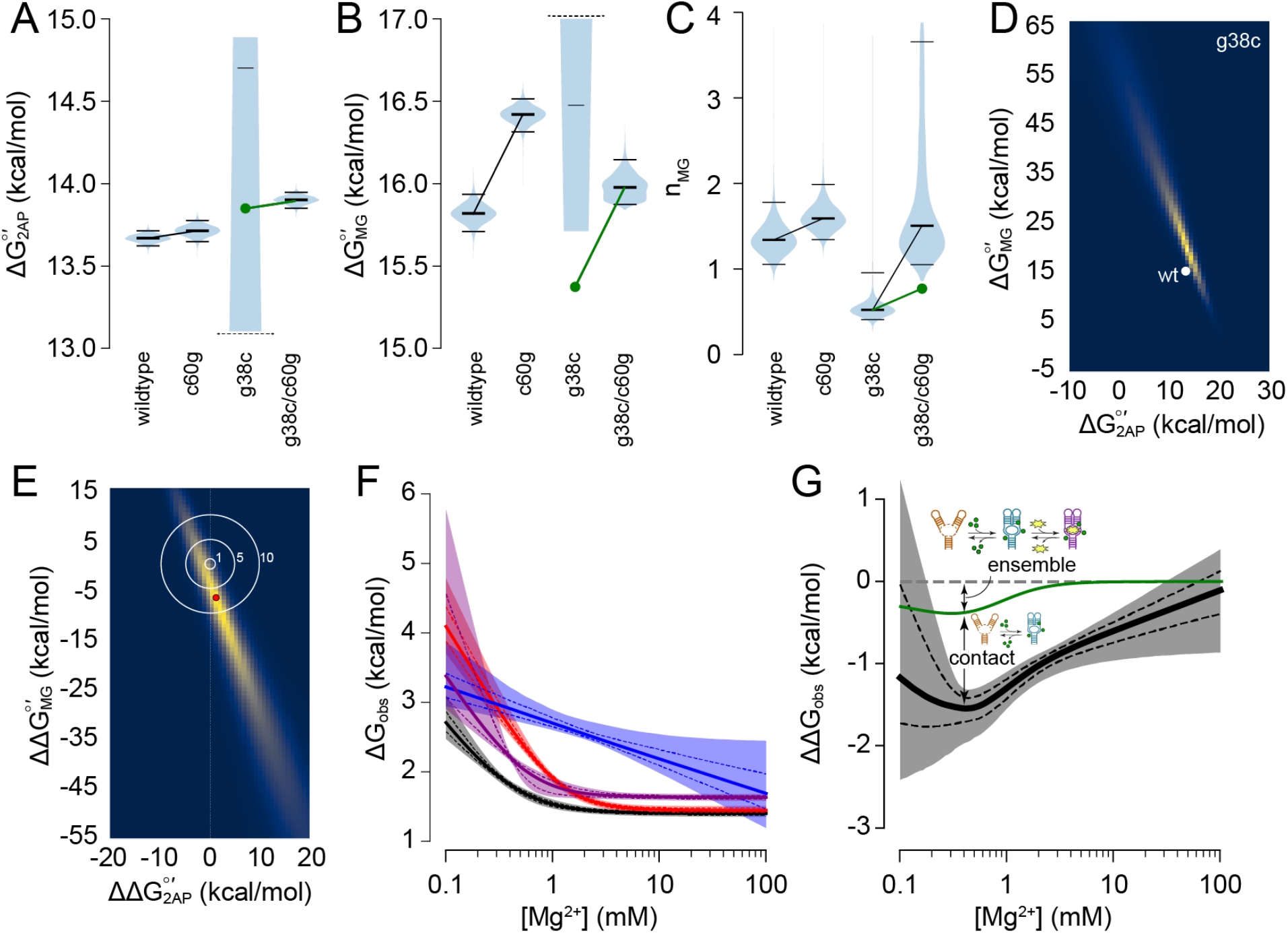
Contact and ensemble epistasis in the g38c/c60g mutant cycle. Panels A-C show marginal distributions of ΔG°’2ap, ΔG°’_MG_ and n_MG_ taken across MCMC samples. The area of the violin plot encompasses all values. The central bar is the median; the top and bottom bars mark the 95% credibility interval. The distributions of ΔG°’_2AP_ and ΔG°’_MG_ for the g38c variant are truncated at the dashed lines. The green lines and points indicate the expected value if the mutations exhibit no epistasis. D) Estimates for ΔG°’_2AP_ and ΔG°’_MG_ across MCMC samples for the g38c variant (dark to light encodes low to high frequency). The median wildtype value is shown as a white circle. E) Estimates of epistasis in ΔG°’_MG_ and ΔG°’_MG_ across MCMC samples. The red point indicates the centroid of the distribution. The concentric circles indicate where the total epistasis summed over ΔG°’_2AP_ and ΔG°’_MG_ is 1, 5, and 10 kcal/mol. F) ΔG_obs_ versus Mg^2+^concentration calculated using Equation 7 for wildtype (black), c60g (red), g38c (blue) and g38c/c60g (purple). Solid lines indicate the median of the MCMC samples; dashed lines indicate the standard deviation; areas indicate the 95% credibility region. G) Epistasis in ΔG_obs_ calculated from the curves in panel F. The green line indicates ΔG_obs_ in the absence of contact epistasis (green points in panels A-C). The small arrows indicate the maximal contributions of ensemble and contact epistasis near 1 mM Mg^2+^.

We calculated contact epistasis in ΔG°’_2AP_, ΔG°’_MG_ and n_MG_ for the g38c/c60g mutant cycle by drawing from each parameter’s MCMC distributions for each variant and applying Equation 9. We observed strong covariation in epistasis between ΔG°’_2AP_ and ΔG°’_MG_ (Fig 4E). The centroid of the distribution yields estimates of the epistasis in each parameter as 1 and −6 kcal/mol, respectively (red point, Fig 4E). Only 0.4% of samples have epistasis < 1 kcal/mol in both ΔG°’_2AP_ and ΔG°’_MG_, strongly suggesting contact epistasis is present in one or both parameters.

To look for ensemble epistasis in the g38c/c60g mutant cycle, we first calculated ΔG_obs_ vs. Mg^2+^ concentration for all four variants (Fig 4F). Because of the strong covariation in our estimates of ΔG°’_2AP_ and ΔG°’_MG_, our confidence in ΔG_obs_ for g38c is high despite low confidence in the absolute magnitudes of these parameters. We found that ΔG_obs_ plateaued for all variants except g38c. This reflects the fact that 2AP binding approaches saturation for all variants except g38c (Fig S3).

We observed striking magnesium-dependent epistasis in ΔG_obs_ for the g38c/c60g mutant cycle (Fig 4G, Equation 10). The value is indistinguishable from zero at low and high Mg^2+^ concentrations. We see epistasis of −1.5 [-1.9, −1.3] kcal/mol, peaking around the point where the riboswitch binds Mg^2+^ and 2AP (Fig S3). The negative sign indicates the double mutant variant is 1.5 kcal/mol *better* at binding 2AP than we would expect based on the effects of the mutations alone. This is driven largely by the deleterious effect of g38c when introduced alone (blue curve; Fig 4F) and aligns with previous measurements of restored 2AP binding upon re-establishing a Watson-Crick-Franklin base pair between these sites in the D conformation (19).

To decompose the interplay between ensemble and contact epistasis for the g38c/c60g mutant cycle, we eliminated contact epistasis from ΔG_obs_ by removing epistasis in ΔG°’_2AP_, ΔG°’_MG_, and n_MG_ as before. For the wildtype, c60g, and g38c/c60g variants, we used the median value for each parameter from its MCMC distribution (Fig 4A-C). To obtain ΔG°’_2AP_ and ΔG°’_MG_ for g38c, we subtracted the effect of c60g on each parameter in the wildtype background from the values for the g38c/c60g double mutant (green points, Fig 4A-B). This removal of contact epistasis led to the solid green curve in Fig 4G, which shows that ensemble redistribution alone contributes up to −0.5 kcal/mol of epistasis at low Mg^2+^ concentrations (labeled arrow, Fig 4G). Contact epistasis further amplifies this effect to yield the maximum epistasis of −1.5 kcal/mol (labeled arrow, Fig 4G). This contact epistasis is driven primarily by epistasis in Mg^2+^ affinity because eliminating epistasis in ΔG°’_MG_ gives a value entirely outside the MCMC samples for g38c (green point; Fig 4B).

### Mutant Cycle #3: c50a/c60g

We next turned to analyzing epistasis in the c50a/c60g mutant cycle. We expected this mutant cycle to yield minimal contact epistasis but significant ensemble epistasis between allosteric sites. These two positions are distant from one another but connect two allosterically linked regions of the protein: the docked helices (c60g) and the adenine binding site (c50a).

When introduced alone, we found that c50a had a deleterious effect of 1.1 [0.9,1.3] kcal/mol on 2AP binding (Fig 5A), but no detectable effect on Mg^2+^ binding (Fig 5B,C). This matches previous observations and makes sense given its proximity to the adenine binding pocket (20). When c50a is combined with c60g, the riboswitch becomes almost unresponsive to Mg^2+^ as measured by 2AP binding. Even at the highest RNA and Mg^2+^ concentrations we studied, we saw only small shifts in 2AP binding (Fig 2B). As a result, the values of ΔG°’_2AP_ and ΔG°’_MG_ for the double mutant are poorly defined.

**Figure 5.**
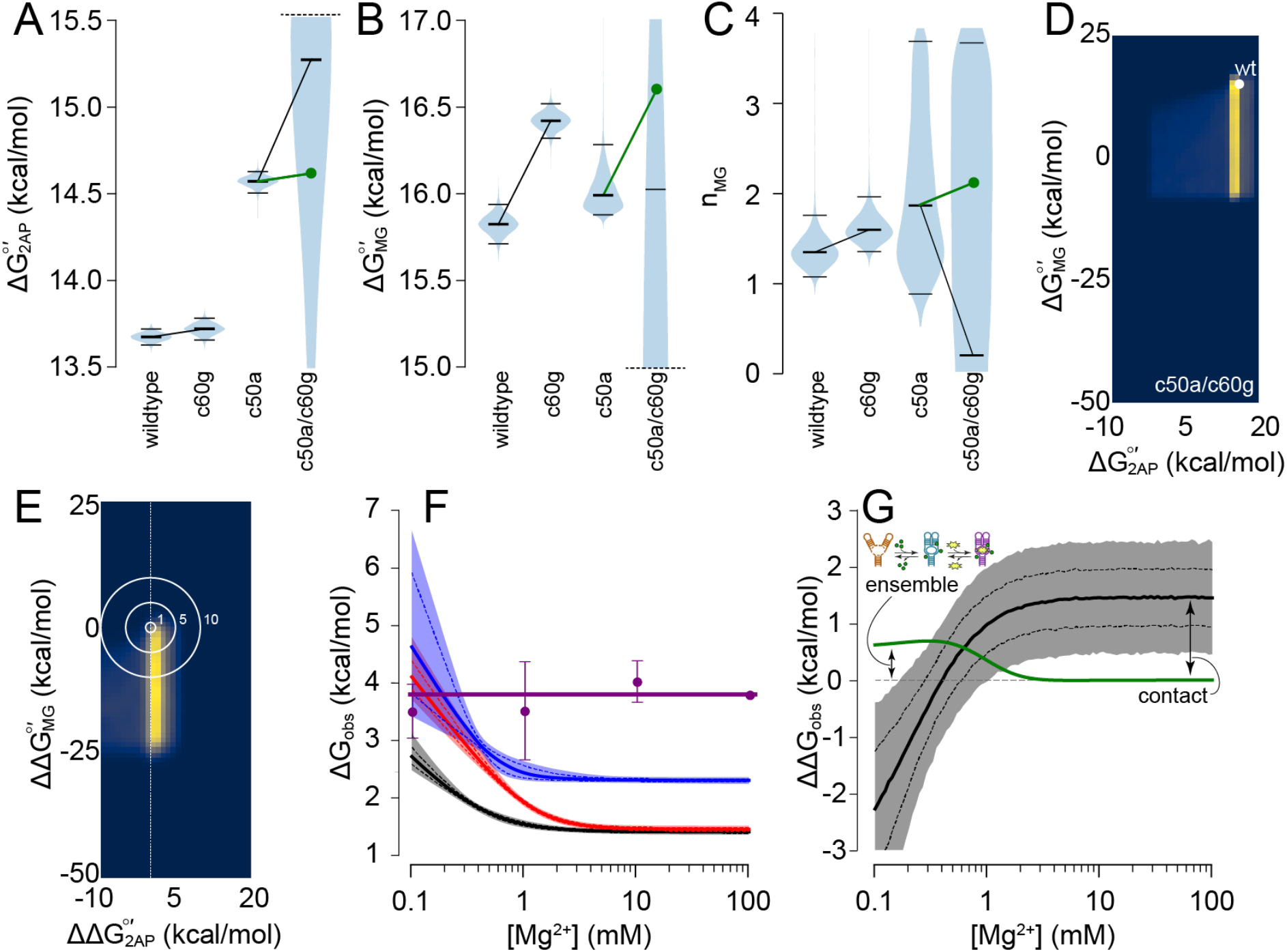
Contact and ensemble epistasis in the c50a/c60g mutant cycle. Panels A-C show marginal distributions of ΔG°’_2AP_, ΔG°’_MG_ and n_MG_ taken from across MCMC samples. The area of the violin plot encompasses all values. The central bar is the median; the top and bottom bars mark the 95% credibility interval. The distributions of ΔG°’_2AP_ and ΔG°’_MG_ for the c50a/c60g variant are truncated at the dashed lines. The green lines and points indicate the expected value if the mutations exhibit no epistasis. D) Estimates of ΔG°’_2AP_ and ΔG°’_MG_ across MCMC samples for the c50a/c60g variant (dark to light encodes low to high frequency). The median wildtype value is shown as a white circle. E) Estimates of epistasis in ΔG°’_2AP_ and ΔG°’_MG_ across MCMC samples. The concentric circles indicate combined epistasis between the two parameters of 1, 5, and 10 kcal/mol. F) ΔG_obs_ versus Mg^2+^ concentration calculated using Equation 7 for wildtype (black), c60g (red), and c50a (blue). Solid lines indicate the median of the MCMC samples; dashed lines indicate the standard deviation; areas indicate the 95% credibility region. The purple points indicate the mean and standard deviation of ΔG_obs_ for c50a/c60g calculated from the binding curves in Fig 2 using Equation 8. G) Epistasis in ΔG_obs_ calculated from the curves in panel F. The green line indicates ΔG_obs_ calculated assuming no contact epistasis (green points in panels A-C). The small arrows indicate the Mg^2+^ concentrations where the epistasis may arise from the ensemble versus contacts.

To investigate contact epistasis for the c50a/c60g mutant cycle we looked at the joint distribution of ΔG°’_2AP_ and ΔG°’_MG_ for the double mutant (Fig 5D). Unlike the previous mutant cycle, we saw no covariation in our estimates of ΔG°’_2AP_ and ΔG°’_MG_: ΔG°’_MG_ is essentially unconstrained by the data. There is some evidence for contact epistasis, even for these poorly defined parameters, as only 1.1% of MCMC samples had ΔG°’_2AP_ and ΔG°’_MG_ that both had epistasis less than 1 kcal/mol. This result is tentative, however, given the poor constraints on parameters given by the experimental data for c50a/g60c.

To look for ensemble epistasis, we next calculated ΔG_obs_ vs. Mg^2+^ concentration for the wildtype, c50a, and c60g variants (Fig 5F). Because the value of ΔG°’_MG_ was unconstrained for the c50a/c60g variant, we could not use Equation 7 to estimate ΔG_obs_. We therefore turned to Equation 8, which allows us to estimate ΔG_obs_ at each Mg^2+^ concentration directly from the 2AP binding curves (Fig 2B). ΔG_obs_ for c50a/c60g estimated using Equation 8 is shown as points in Fig 5F. We observed no apparent change in ΔG_obs_ for c50a/c60g as a function of Mg^2+^ concentration, reflecting its unresponsiveness to Mg^2+^ ions. To calculate epistasis in ΔG_obs_ using Equation 10, we used the average value of ΔG_obs_ for c50a/c60g across all measured Mg^2+^ concentrations (3.8 ± 0.5 kcal/mol; solid purple line, Fig 5F).

We observed magnesium-dependent epistasis in ΔG_obs_ for the c50a/c60g mutant cycle (Fig 5G): the value starts negative, then saturates just above 1 mM Mg^2+^. At low concentrations, the epistasis arises from the difference in the responsiveness of the variants to Mg^2+^. At high Mg^2+^concentrations, the flat epistasis curve arises because the four variants reach different final populations of D·A (Fig S3). The epistasis is positive because the double mutant is worse at binding 2AP than would be expected based on the individual effects of the mutations.

We next investigated the interplay between contact and ensemble epistasis in the c50a/c60g mutant cycle. As before, we calculated epistasis in ΔG_obs_ after removing epistasis in ΔG°’_2AP_, ΔG°’_MG_ and n_MG_ for all variants. For this analysis, we used the median values of our estimates for wildtype, c50a, and c60g variants and the green points for c50a/c60g (Fig 5A-C). The resulting ensemble epistasis in ΔG_obs_ is shown as a green line in Fig 5G. For this mutant cycle, the ensemble contributes a maximum of 0.7 kcal/mol of epistasis at Mg^2+^ concentrations below 1 mM (labeled arrow, Fig 5G). At higher Mg^2+^ concentrations, contact epistasis dominates (labeled arrow, Fig 5G). Given our uncertainty in the thermodynamic parameters, we cannot resolve what contact(s) and corresponding magnitudes lead to the observed epistasis.

### General patterns of concentration-dependence

We next asked if there were general rules that might allow one to make hypotheses about the source of epistasis given an experimental observable, even without detailed information about the thermodynamic ensemble under study. The riboswitch ensemble provides a useful framework for posing this question. We can rewrite our energetic observable as follows, pulling out the 2AP binding constant:

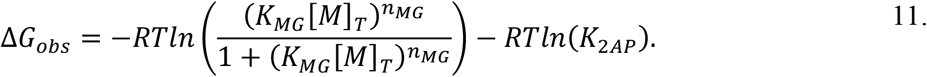

When we do so, we see that ΔG_obs_ reduces to a Hill binding model (left term) with a Mg^2+^-independent offset controlled by 2AP affinity (right term). Such a model is generic and applicable to many systems. We can therefore think about the interplay of contact and ensemble epistasis in the adenine riboswitch as a model for many macromolecules of interest. What patterns arise if we introduce contact epistasis in K_2AP_ (target affinity), K_MG_ (allosteric effector affinity) and n_MG_ (allosteric effector cooperativity)?

In Figure 6, we calculated epistasis in ΔG_obs_ for otherwise non-epistatic mutant cycles in which we injected varying amounts of epistasis into these equilibrium constants. We observe that each equilibrium constant gives a distinct effector-dependent epistatic signal. First, epistasis in 2AP affinity has no concentration dependence: it acts as a global offset to ΔG_obs_ (Fig 6A). We observed this in our experimental data: epistasis in ΔG°’_2AP_ led to a global offset (Fig 3E). The affinity for 2AP is independent of the population of the 2AP binding-competent conformation and thus does not depend on Mg^2+^ concentration.

**Figure 6.**
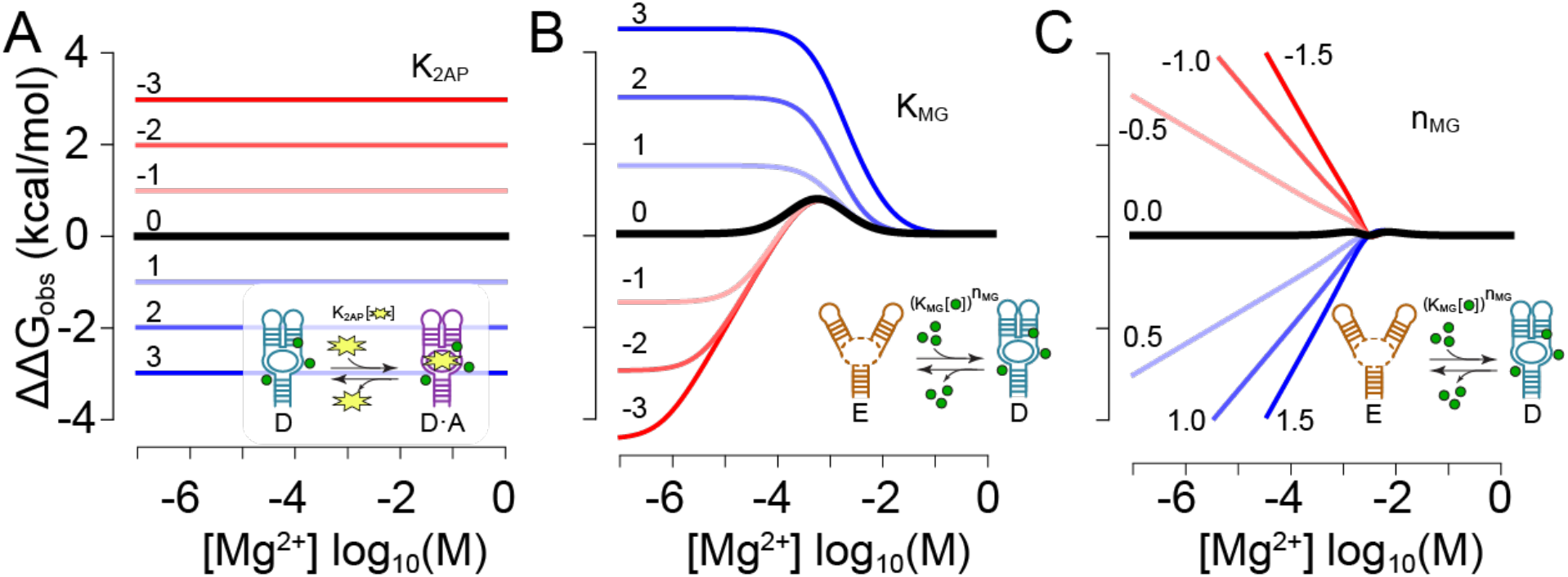
Contact epistasis alters epistasis in ΔG_obs_. Panels A-C show calculated effects of contact epistasis in K_2AP_ (A), K_MG_ (B), or n_MG_ (C) on epistasis in ΔG_obs_ vs. Mg^2+^ concentration. The black curves show epistasis in ΔG_obs_ for two mutations that have no contact epistasis. Mutations were modeled as having an effect of +1 kcal/mol (A and B) or +0.5 ions (C). The colored curves are epistasis in ΔG_obs_ given the positive (blue) or negative (red) contact epistasis indicated next to each curve. Cartoon insets indicate which equilibrium is being perturbed.

Contact epistasis in Mg^2+^ affinity gives more complex patterns. In the absence of contact epistasis in K_MG_, we observe a peak in epistasis (black curve, Fig 6B; Fig 1C). The peak arises from the redistribution of conformations within the ensemble: there is no contact epistasis anywhere in the system. As we add contact epistasis, it modulates the signal for ensemble epistasis in a nonlinear fashion: the contact epistasis itself redistributes the ensemble, amplifying the ensemble epistasis. The largest effects of epistasis in K_MG_ occur at low Mg^2+^ concentrations where a change in Mg^2+^binding can alter the relative population of the D·A conformation, and thus ΔG_obs_ (Fig S3). This epistasis is lost at high Mg^2+^ because the molecule is saturated with Mg^2+^ and 2AP binding now controls ΔG_obs_ (Fig S3). We observed this pattern for the g38c/c60g mutant cycle, where strong contact epistasis in ΔG°’_MG_ amplified ensemble epistasis at low and moderate Mg^2+^concentrations, then dropped to zero at high Mg^2+^ concentrations (Fig 4G).

Finally, epistasis in ion uptake leads to a different pattern of magnesium-dependent epistasis. As with epistasis in K_MG_, contact epistasis in n_MG_ plays a role at low Mg^2+^ concentration, then goes to zero when other factors outside Mg^2+^ binding are the most important determinants of the energetics of the system. Because the ion uptake coefficient controls the sensitivity to Mg^2+^— mathematically, it is equivalent to a Hill coefficient (31)—contact epistasis in n_MG_ manifests as a linear dependence between the log of Mg^2+^ concentration and epistasis. We saw no evidence for this particular form of epistasis in our riboswitch mutant cycles.

This rogues’ gallery of epistatic effects provides a useful lens to generate hypotheses about the origins of epistasis in a macromolecule, even without a detailed model of the thermodynamic ensemble for that system. If one were to observe a fixed epistatic signal across multiple concentrations of an allosteric effector, it would suggest contact epistasis on some feature of the macromolecule that does not depend on the effector concentration (Fig 6A). If one observed a peak in epistasis that dropped to zero at both high and low effector concentrations—concentration regimes in which few conformations are populated—the pattern would suggest ensemble epistasis and minimal contact epistasis (Fig 6B). If one observed high epistasis at low effector concentration that dropped to zero at high effector concentration, it would suggest contact epistasis in the parameters controlling effector binding (Fig 6B). And, finally, if one saw a monotonic change in epistasis with the logarithm of effector concentration, it would suggest contact epistasis in binding cooperativity (Fig 6C).

These rules of thumb are not definitive and require system-specific validation (to say nothing of the complexity that could arise if multiple classes of contact epistasis contribute to the observed signal). However, they do provide a useful starting point. One might imagine a high-throughput experiment conducted at a few allosteric effector concentrations. A simple analysis of the effector-dependent epistasis in the system would provide initial hints about the energetic couplings within the system. This could set up deeper biophysical investigation, possibly leading to fitting a thermodynamic model to the system (as demonstrated in this work) to extract the relevant thermodynamic parameters and epistatic mechanisms (16, 17).

## Discussion

We set out to investigate the relative contributions of contact and ensemble epistasis between mutations in the adenine riboswitch. We found evidence for both forms of epistasis in all three mutant cycles. Our work reveals that ensemble epistasis—previously measured in proteins (17)—also occurs in RNA and is likely a generic feature of allosteric macromolecules. Further, we found that contact and ensemble epistasis interact in a nonlinear fashion. Even when the magnitude of ensemble epistasis was low (~0.5 kcal/mol) in one mutant cycle, the ensemble could strongly attenuate the magnitude of contact epistasis (from ~6 kcal/mol to ~1.5 kcal/mol, Fig 4E,G). Thus, understanding intramolecular epistasis in this molecule requires accounting for both contact and ensemble mediated effects.

### Interplay between ensemble and contact epistasis

In our previous theoretical and experimental work, we observed that mutations led to high-magnitude epistasis in thermodynamic observables defined by the ensemble (14, 17), even without any epistasis between mutations at the level of equilibrium constants. This was *not* the case for adenine riboswitch. If we removed epistasis from equilibrium constants (ΔG°’_2AP_, ΔG°’_MG_, and n_MG_), ensemble epistasis in ΔG_obs_ hovered around kT (0.6 kcal/mol). This can be seen in the green curves shown in Fig 3E, 4G, and 5G, which have peak magnitudes of 0.3 (a35u/c60g), −0.5 (g38c/c60g), and 0.7 (c50a/c60g) kcal/mol, respectively.

When we included contact epistasis, however, we observed much larger concentrationdependent epistasis in ΔG_obs_ for two of the three cycles: g38c/c60g (−1.5 kcal/mol), and c50a/c60g (1.5 kcal/mol). Concentration-dependent epistasis folds together contact and ensemble mechanisms in a nonlinear fashion. This is seen most clearly for g38c/c60g. This cycle had strong contact epistasis because it inverted the polarity of a Watson-Crick-Franklin G-C base pair. Contact epistasis in either ΔG°’_2AP_ or ΔG°’_MG_ is necessary to explain the observed binding curves (Fig 4B, Fig 4C). Although we could not unambiguously determine which parameter exhibited epistasis due to strong covariance, the median estimate of contact epistasis was −6 kcal/mol in ΔG°’_MG_ and 1 kcal/mol in ΔG°’_2AP_ (red point, Fig 4E). This result is plausible, as the above value of ΔG°’_MG_ is consistent with a previous measurement of the g38/c60 interaction energy (19), as well as the expected energy for a G-C base pair (37). Further, we would anticipate that epistasis from an inverted G-C base pair in the docked-loop conformations would manifest largely in ΔG°’_MG_ rather than indirectly in ΔG°’_2AP_ (Fig 1B).

Despite this strong contact epistasis (−6 kcal/mol) in the g38c/c60g mutant cycle, we observed epistasis of only −1.5 kcal/mol in our ensemble observable (ΔG_obs_). The riboswitch’s conformational ensemble attenuates the strong contact epistasis, making epistasis in the observable much lower than the epistasis in the underlying equilibrium constant. We note that even MCMC samples with contact epistasis up to −45 kcal/mol in ΔG°’_MG_ and 10 kcal/mol in ΔG°’_2AP_(distribution, Fig 4E) were accommodated with only ~1.5 kcal/mol epistasis in ΔG_obs_ (gray area, Fig 4G). Although these massive energetic effects on ΔG°’_MG_ and ΔG°’_2AP_ are chemically implausible, they illustrate an important point. Epistasis in ΔG_obs_ arises due to shifts in the relative population of ensemble conformations—E, D, and D·A—rather than directly from the energetic effects of mutations on each conformation. Massive effects on the energy of each conformation due to physical contacts can be subsumed by the ensemble if these effects do not meaningfully redistribute the probabilities of the conformations.

### The importance of multistate design

Our work points to the fundamental importance of conformational landscapes in modulating the effects of mutations and epistasis between them. Efforts to engineer or design macromolecules by focusing on individual conformations may face unexpected results if the macromolecule populates multiple conformations in solution.

As a hypothetical example, consider an attempt to optimize the adenine riboswitch’s ligand-bound conformation for stronger ligand binding affinity. Perhaps a pair of mutations are introduced to form another Watson-Crick-Franklin base pair to bridge the docked-loop interface. Structure-based calculations on the ligand-bound conformation might predict a certain favorable energetic interaction between the two mutations. They might incorrectly calculate the energetic effect on ligand binding, however, by failing to examine how the new base pair reshapes the conformational landscape. These considerations point to the value of multi-state design to better account for the interplay between contact and ensemble sources of epistasis (38, 39), especially since many biological macromolecules must toggle between multiple conformational states under physiological conditions for function.

### Implications for allostery

This work also provides an intriguing lens for attempting to understand the molecular basis of allostery. Epistasis and allostery between sites have similar energetic origins: introduction of a new stabilizing or destabilizing interaction at one site alters the activity at another. Despite this basic equivalence, allostery and epistasis differ in their concentration-dependence. Mutations are a fixed perturbation, independent of environment. In contrast, allosteric interactions depend on the concentrations of allosteric effectors. By combining measurements of epistasis (fixed perturbation to the probability of a conformation) with an allosteric perturbation (addition of allosteric effector) we gain insight into the thermodynamic basis for communication between sites in a macromolecule. In particular, we can separate epistatic effects into “contact” and “ensemble” categories, which may allow us to disentangle communication occurring *through* a molecule (13) from communication occurring through a redistribution of conformational states (15).

We see a hint of long-distance communication through a conformation rather than the ensemble for c50a/c60g. For this mutant cycle, the vast majority of MCMC samples exhibited epistasis in ΔG°’_MG_ (Fig 5E), even though these two mutations are ~34 Å apart in the structure. One way to view this result is that communication is occurring via one of the conformations in the ensemble, rather than through redistributing the ensemble. The thermodynamic parameters for this cycle were poorly determined, so this observation remains tentative. We bring it up here, however, as an example of a promising way to disentangle through-conformation from ensemble-based mechanisms of long-distance communication in macromolecules: look for evidence of epistasis between distant mutations in the microscopic equilibria that define the ensemble (e.g., ΔG°’_MG_)versus epistasis in the overall thermodynamic observable for the system (e.g., ΔG_obs_).

### Identifying sources of epistasis

Finally, our work suggests a few “rules of thumb” that researchers could use to make hypotheses about the roles of contact and ensemble mechanisms in a macromolecule of interest, even given limited measurements. This is particularly apropos for high-throughput studies, where one might be able to measure binding, or some other thermodynamic observable, as a function of allosteric effector. One might imagine using these rules to classify sites for further study, or to help develop a more rigorous mechanistic model of an ensemble for further characterization.

We observed a few basic patterns in our experimental (Figs 3-5) and model studies (Fig 6). First, contact epistasis in the observed conformation (D·A, for the riboswitch) gives rise to epistasis independent of effector concentration (Fig 6A). Second, ensemble epistasis with no contact epistasis yields a peak in epistasis centered at the allosteric effector concentration that maximizes ensemble diversity (black line, Fig 6B) (14). Third, a convolution of contact and ensemble epistasis appears as a peak in epistasis, but with non-zero epistasis at high or low effector concentration (colored lines, Fig 6B). Fourth, contact epistasis in cooperativity (i.e., the ion uptake coefficient or a Hill coefficient) appears as epistasis that changes linearly as with the log of the allosteric effector concentration (Fig 6C). Intriguingly, we also observed very similar patterns of epistasis versus allosteric effector in our previous study of ensemble epistasis in the lac repressor (17).

Further work will be needed to generalize these results, with a particular focus on understanding how these patterns would change if several classes of contact epistasis were in play. It will also be important to explore different sorts of ensembles, with different numbers of conformations and linkages between allosteric effector and observable. This said, given the generic form of the mathematical model of the riboswitch ensemble (Equation 11), we expect these findings to generalize well. Overall, our work highlights the importance of considering effects of mutations on multiple conformations to understand epistasis and suggests a few rules-of-thumb for disentangling contact and ensemble epistasis in macromolecules.

## Acknowledgments

We would like to thank members of the Harms lab, past and present, for their invaluable input. This work was funded by a National Science Foundation CAREER Award DEB-1844963 (M.J.H.).

---

**Table S1:** AIC test comparing models of differing complexity.

| Model (# states, # parameters) | AIC value | AIC probability |
| --- | --- | --- |
| 3.3 | -1033.26 | 0.616 |
| 3.4 | -1031.26 | 0.226 |
| 4.4 | -1030.54 | 0.158 |
| 4.5 | -996.13 | $8.654 \times 10^{-9}$ |
| 2.3 | -630.57 | $3.615 \times 10^{-88}$ |

**Table S2:** Characteristics of marginal MCMC distributions for 3-state, 3-parameter model. Values shown are the median, 2.5% (bottom) and 97.5% (top) values from each distribution.

| genotype | parameter | median | bottom | top |
| --- | --- | --- | --- | --- |
| wt | $\ln(K_{2AP})$ | -2.4 | -2.4 | -2.3 |
| wt | $\ln(K_{MG})$ | -6.0 | -6.2 | -5.8 |
| wt | $n_{mg}$ | 1.5 | 1.1 | 2.1 |
| a35u | $\ln(K_{2AP})$ | -2.3 | -2.4 | -2.2 |
| a35u | $\ln(K_{MG})$ | -6.6 | -6.9 | -6.2 |
| a35u | $n_{mg}$ | 1.6 | 1.2 | 4.5 |
| g38c | $\ln(K_{2AP})$ | 14.5 | -4.1 | 110.3 |
| g38c | $\ln(K_{MG})$ | -57.7 | -289.5 | -7.0 |
| g38c | $n_{mg}$ | 0.4 | 0.2 | 1.0 |
| c50a | $\ln(K_{2AP})$ | -3.9 | -4.0 | -3.8 |
| c50a | $\ln(K_{MG})$ | -6.3 | -6.8 | -6.1 |
| c50a | $n_{mg}$ | 2.3 | 0.9 | 4.8 |
| c60g | $\ln(K_{2AP})$ | -2.4 | -2.5 | -2.3 |
| c60g | $\ln(K_{MG})$ | -7.0 | -7.2 | -6.8 |
| c60g | $n_{mg}$ | 1.9 | 1.5 | 2.4 |
| a35u/c60g | $\ln(K_{2AP})$ | -2.9 | -3.0 | -2.7 |
| a35u/c60g | $\ln(K_{MG})$ | -7.3 | -7.7 | -7.1 |
| a35u/c60g | $n_{mg}$ | 1.5 | 1.1 | 4.2 |
| g38c/c60g | $\ln(K_{2AP})$ | -2.8 | -2.8 | -2.7 |
| g38c/c60g | $\ln(K_{MG})$ | -6.3 | -6.6 | -6.1 |
| g38c/c60g | $n_{mg}$ | 1.7 | 1.1 | 4.7 |
| c50a/c60g | $\ln(K_{2AP})$ | -5.1 | -11.4 | 21.9 |
| c50a/c60g | $\ln(K_{MG})$ | 14.8 | -6.3 | 33.0 |
| c50a/c60g | $n_{mg}$ | -0.1 | -4.8 | 4.8 |

**Fig S1:**
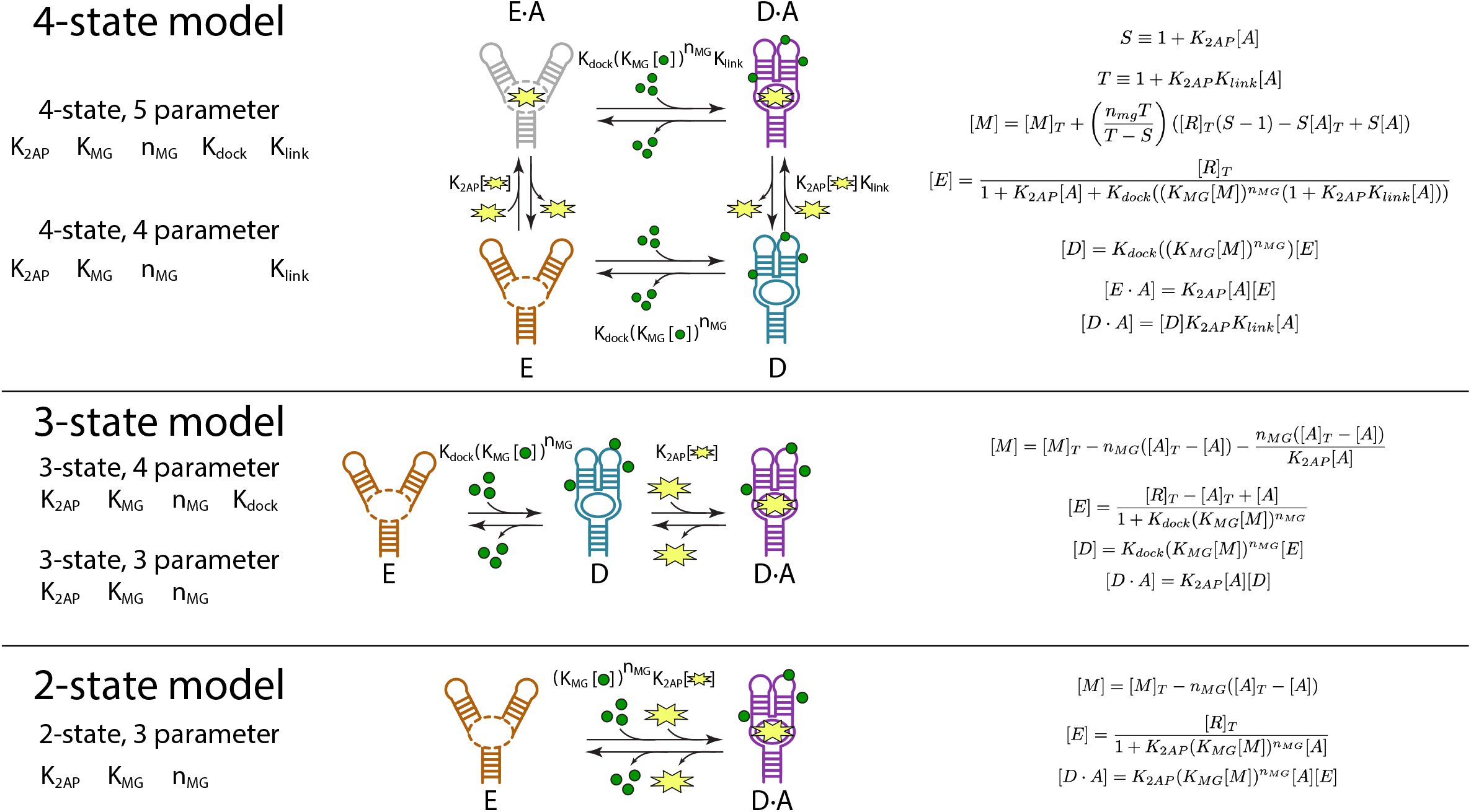
Variants of the adenine riboswitch thermodynamic model. Parameters required are shown on the left; model schema are shown in the middle; mathematical expressions for the the different species are shown on the right. The colors and iconography in the schema match those shown in Figure 1A. In the mathematical expressions, [M] and [A] indicate the free Mg^2+^ and 2AP concentrations; [M]_T_ and [A]_T_ indicate their total concentrations. [R]_T_ is the total RNA concentration. To estimate species concentrations, We numerically found values for [A] that satisfied the conservation of mass in RNA, Mg^2+^, and 2AP.

**Fig S2.**
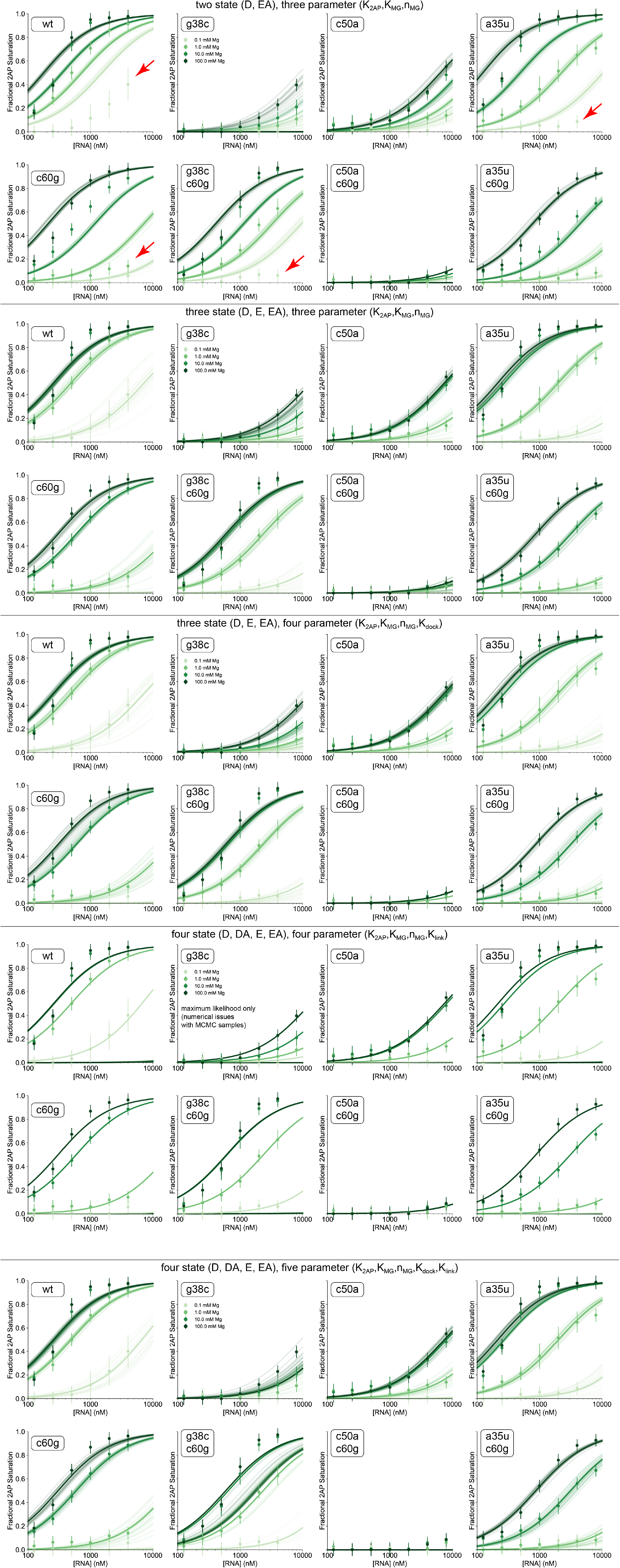
Fits for different binding models to 2AP binding data for each mutant. Model fits are arranged from less complex to more complex, top to bottom. The model and included parameters are shown above each block of eight plots (See Fig S1 for details on each model).Each panel shows the fraction of 2AP bound, measured by relative 2AP fluorescence compared to free and fully quenched controls, as a function of riboswitch concentration. The green shade indicates the total Mg2+ concentration, labeled on the plots. Each point is the average of at least three experimental replicates, with standard deviations indicated as error bars. Lines were calculated using parameters taken from 50 randomly selected MCMC samples. Systematic deviations between the model and experimental data are indicated with red arrows. For the four state, four parameter data, the model was numerically unstable. The maximum likelihood fits but not MCMC samples are shown.

**Fig S3.**
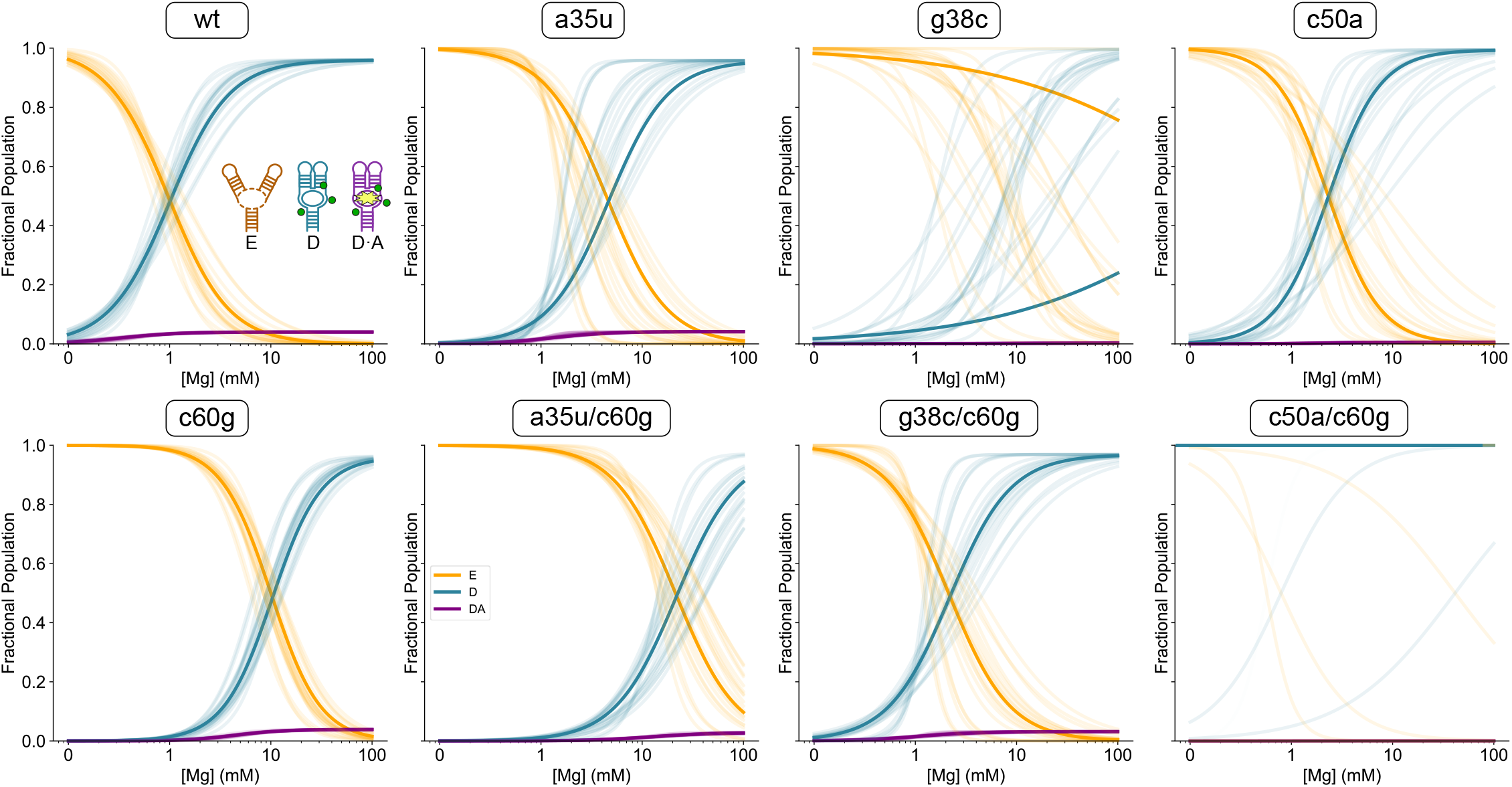
Magnesium-dependent changes in the populations of conformations in the ensemble. Each panel shows the population of E (orange), D (teal), and D·A (purple) as a function of total [Mg^2+^] for the indicated RNA variant. Faded lines were calculated using parameters taken from 50 randomly selected MCMC samples of the three-state, three-parameter model shown in Fig 1A. The solid lines are the median populations across MCMC repli cates.

## Notes

### Competing Interest Statement

The authors have declared no competing interest.

